# Single-cell whole-genome sequencing reveals the functional landscape of somatic mutations in B lymphocytes across the human lifespan

**DOI:** 10.1101/535906

**Authors:** Lei Zhang, Xiao Dong, Moonsook Lee, Alexander Y. Maslov, Tao Wang, Jan Vijg

## Abstract

The accumulation of mutations in somatic cells have been implicated as a cause of ageing since the 1950s^1,2^. Yet, attempts to establish a causal relationship between somatic mutations and ageing have been constrained by the lack of methods to directly identify mutational events in primary human tissues. Here we provide detailed, genome-wide mutation frequencies and spectra of human B lymphocytes from healthy individuals across the entire human lifespan, from newborns to centenarians, using a recently developed, highly accurate single-cell whole-genome sequencing method^3^. We found that the number of somatic mutations increases from <500 per cell in newborns to >3,000 per cell in centenarians. We discovered mutational hotspot regions, some of which, as expected, located at immunoglobulin genes associated with somatic hypermutation. B cell-specific mutation signatures were observed associated with development, ageing or somatic hypermutation (SHM). The SHM signature strongly correlated with the signature found in human chronic lymphocytic leukemia and malignant B-cell lymphomas^4^, indicating that even in B cells of healthy individuals the potential cancer-causing events are already present. We also identified multiple mutations in sequence features relevant to cellular function, i.e., transcribed genes and gene regulatory regions. Such mutations increased significantly during ageing, but only at approximately half the rate of the genome average, indicating selection against mutations that impact B cell function. This first full characterization of the landscape of somatic mutations in human B lymphocytes indicates that spontaneous somatic mutations accumulating with age can be deleterious and may contribute to both the increased risk for leukemia and the functional decline of B lymphocytes in the elderly.

## Main

To accurately detect a full complement of mutations in somatic cells is a major technical challenge because of the random nature and very low abundance of most somatic mutations. This means that sequencing DNA from cell populations will show the germline genotype rather than *de nov*o somatic mutations^5^. The solution to this problem, i.e., single-cell sequencing, is hampered by the high error rate of genome amplification procedures required for single cell genomics^6-8^. We recently developed a highly accurate, single cell multiple displacement amplification (SCMDA) procedure to comprehensively determine the full spectrum of base substitutions in a single somatic cell and validated the procedure by comparing mutations identified from SCMDA-amplified single cells with unamplified DNA from clones of the same cell population^3^.

Here, we used SCMDA to assess mutation accumulation as a function of age in human B lymphocytes from healthy individuals varying in age from 0 (newborns) to over 100 years (centenarians). Mutation accumulation in these cells could be a causal factor in both the age-related increase in the incidence of B cell leukemia^9^ and the observed age-related functional decline, such as loss of B cell activation^10^. We sequenced the genome of 56 single B lymphocytes and that of their corresponding 14 bulk DNAs using the remaining PBMCs (Methods; Supplementary Figs. 1, 2). The average sequencing depth across the genome was 27.9±2.9 (s.d.) and 25.3±2.3 for the single cells and bulk DNAs, respectively. For single cells 51.4±4.7 % of the genome was covered at a depth of ≥ 20X; For bulk DNAs this was 78.6±5.1 % (Supplementary Table 1; Methods). Somatic SNVs in single cells were identified at a depth of ≥ 20X using SCcaller, a software tool developed specifically for single-cell SNV detection by filtering out potential artifacts due to whole-genome amplification (Methods)^3^. Germline variants were filtered out using the bulk DNAs of the same donors. Using Sanger sequencing, we confirmed 62 of 65 (95.4%) mutations randomly selected, indicating the high specificity of the variant calls (Supplementary Fig. 3 and Supplementary Table 2). Because all somatic SNVs should be heterozygous due to the extremely low chance that mutations occur at the exact same site on both alleles, heterozygous germline SNP calling in the same cell is a good control for measuring sensitivity, which appeared to be on average 60% (Supplementary Table 3; Methods). That is, 60% of the heterozygous SNPs identified in bulk DNA were also identified in a single cell. The less than 100% sensitivity is a consequence of the stringent filtering, which avoids false positives at the cost of occasionally missing a true variant.

After adjusting for genomic coverage and sensitivity, the numbers of SNVs per cell were estimated as between 237 and 5,862, with one extreme outlier of 11,765 (Figs. 1a, Supplementary Table 3). The median number of mutations per cell was found to increase significantly with the age of the donor (*P*=3.93×10^−4^, Fig. 1a), with on average 463.4 in the newborns (n=8 cells from two donors), 1,181.9 in the 27-30 year olds (n=8 cells from two donors), 2,101.7 in the 52-75 year olds (n=24 from 6 donors) and 3,127.0 in the 97-106 year olds (n=16 from 4 donors).

**Fig. 1.**
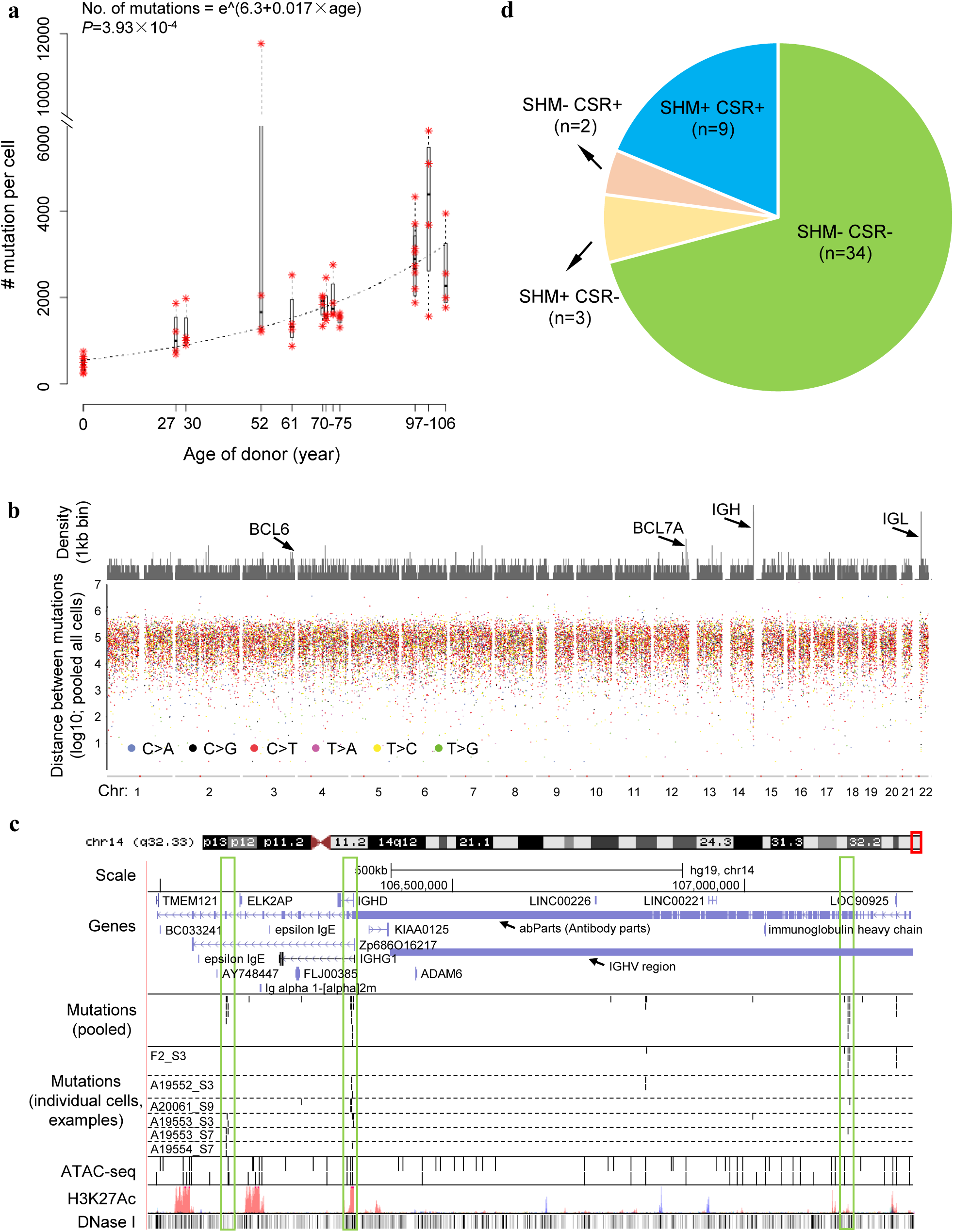
Somatic mutations accumulate with age in human B lymphocytes. **a.** The number of somatic mutations per genome as a function of age. Red asterisks indicate individual cells. Each box corresponds to summary statistics of four cells of one donor. The regression was performed on the median numbers of mutations of the 4 cells from each donor, using exponential model non-linear least squares regression. **b.** Rainfall plot illustrates distances of neighboring mutations. The top panel shows the density of mutations (56 cells pooled) in kb bins. The bottom panel shows distances of each mutation to its closest other mutation. **c.** Mutation hotspots in IGH regions visualized using the UCSC genome browser. In the mutation panels, each bar represents one SNV. In the ATAC-Seq panel, obtained from pooled B cells from one young and one old individual, each bar represents one open chromatin peak. Other panels were obtained from annotations of the UCSC genome browser. The three vertical boxes highlight three hotspots identified in this region. **d.** The SHM and CSR status of the 48 adult B cells (all 8 B cells from cord blood are SHM-CSR-). The majority of adult B cells are as expected classified into two categories, i.e., SHM-CSR-(naïve B cells) and SHM+CSR+ (memory B cells). A few cases of SHM+CSR- and SHM-CSR+ cells are known as non-switched memory B cells and GC-independent memory B cells, respectively.

While virtually all mutations were unique for each cell, a total of 24 recurrent mutations, i.e., shared between cells, were identified. These recurrent mutations were only present in cells from the same donor, which was validated by Sanger sequencing of the recurrent mutations in all samples (Supplementary Fig. 4 and Supplementary Table 2). The cells that share mutations most likely derive from the same ancestor hematopoietic stem cell in which the mutation originally occurred. Of note, the recurrent mutations were mostly found in elderly donors (*P*=0.0115, permutation test, one-tailed, 20,000 permutations), which is in keeping with the observed decrease in the number of active lineages of hematopoietic stem cells during ageing^11^. However, the numbers of recurrent mutations were small and none occurred in the cancer driver genes used to identify clonal hematopoiesis of indeterminate potential (CHIP)^12^; we also applied the same method as described in ref^12^ and found that none of our 14 donors displayed CHIP (Methods).

While the observed increase in mutation frequency with age is in keeping with previous data on other cell types in both humans and mice^13-17^, accurate numbers obtained by single-cell sequencing are sparse and have not been thoroughly validated^6^. In terms of validation, B lymphocytes offer the advantage of endogenous control loci in the form of mutational hotspots at immunoglobulin genes^18^. After pooling all mutations of the 56 cells the large majority of mutations distributed randomly across the genome (Supplementary Fig. 5a-c). However, we noticed already at Mb level a substantial enrichment of mutations at one end of chromosome 14 (Supplementary Fig. 5b,c), which harbors immunoglobulin gene sequences^18^. At the kilobase (kb) level, we identified 24 mutational hotspots, defined as regions with multiple mutations separated by less than 5 kb from their nearest neighbor (using the “shower” algorithm; Fig. 1b, Supplementary Table 4; Methods)^19-21^. The 24 mutational hotspots are composed of 4 to 12 mutations in regions of 469 to 8,970 bp corresponding to a mutation frequency of 2.79×10^−3^ per bp per cell in agreement with previous findings as determined from cloned, PCR-amplified target regions^22,23^; the average genome-wide mutation frequency is only about 10^−7^ per bp.

Of the 24 hotspots (Supplementary Fig. 6), only 5 were located in or close to immunoglobulin gene regions, i.e., 3 hotspots in the immunoglobulin H chain (IGH) region on chromosome 14 (Fig. 1c), one in the immunoglobulin L (IGL) chain region and one other downstream of this region on chromosome 22 (Supplementary Fig. 7). The other 19 hotspots are likely off-target effects of activation-induced cytidine deaminase (AID)^18^, the enzyme that initiates somatic hypermutation (SHM). Some of these hotspots have been described previously by others, such as *BCL6* (B-cell CLL/lymphoma 6)^24^ and *BCL7A* (BCL tumor suppressor 7A)^25^. To our knowledge, nobody has as yet demonstrated B lymphocyte mutational hotspots collectively across a single genome. However, when comparing to a mouse AID (mAID) ChIP sequencing dataset^26^, we found substantial overlap of the hotspot regions we identified with genes in the mAID dataset (odds ratio=1.93; Supplementary Fig. 8). While not significant (*P*=0.134, one tailed Fisher’s exact test) this finding contributes to the notion that the hotspots we found are AID off-targets. Indeed, this conclusion is strengthened by the fact that about half of the 24 hotspots have been reported in other studies as associated with human lymphoma, leukemia, or AID off-target loci in mouse B cells^24,25,27-31^ (Supplementary Table 4). Moreover, the hotspots significantly overlap with or are close to open chromatin regions, transcription start sites (TSS), and transcription factor (TF) binding sites associated with AID binding (Supplementary Table 4; Methods).

Based on the presence or absence of mutations in the 5 immunoglobulin hotspots we distinguished SHM^+^ from SHM^−^ cells. In addition to SHM, we also identified class switch recombination (CSR) in some B cells, allowing to distinguish memory B cells (SHM^+^ and CSR^+^) from the SHM^−^/CSR^−^, navïe B cells (Fig. 1d, Supplementary Fig. 9; Methods). Of note, the SHM^+^ cells had a significantly higher mutation frequency than the SHM^−^ cells (*P*=5.19×10^−5^, Supplementary Fig. 10a). A significant age-related increase of mutations in both cell types was observed when analyzing all cells together with both ageing and SHM as variables (*P*=3.07×10^−4^ for ageing, Supplementary Fig. 10a). An age-related increase is also statistically significant when testing SHM^−^ cells alone, with ageing as the only variable (*P*=1.08×10^−4^, Supplementary Fig. 10b).

Next, we analyzed the mutation spectra. In addition to the affected base (Supplementary Fig. 11a, b; Methods), we also included their two flanking bases in the analysis. This allowed us to comparatively analyze mutation spectra as a function of age and SHM status. The results were compared to the spectrum we previously obtained for human fibroblasts^3^ (Supplementary Fig. 11c). By using non-negative matrix factorization (NMF)^20,32^, we decomposed the mutation spectra into mutation signatures, i.e., signatures A to D (Figs. 2a, b; Methods). Based on these signatures, we were able to, first, show that somatic mutations in B lymphocytes (Signatures A, B and D) are substantially different from those in fibroblasts (Signature C). Second, we were able to identify mutations specifically associated with development (Signature A), ageing (Signature B; *P*=9.25×10^−4^ with age, Supplementary Fig. 12) and SHM (Signature D).

**Fig. 2.**
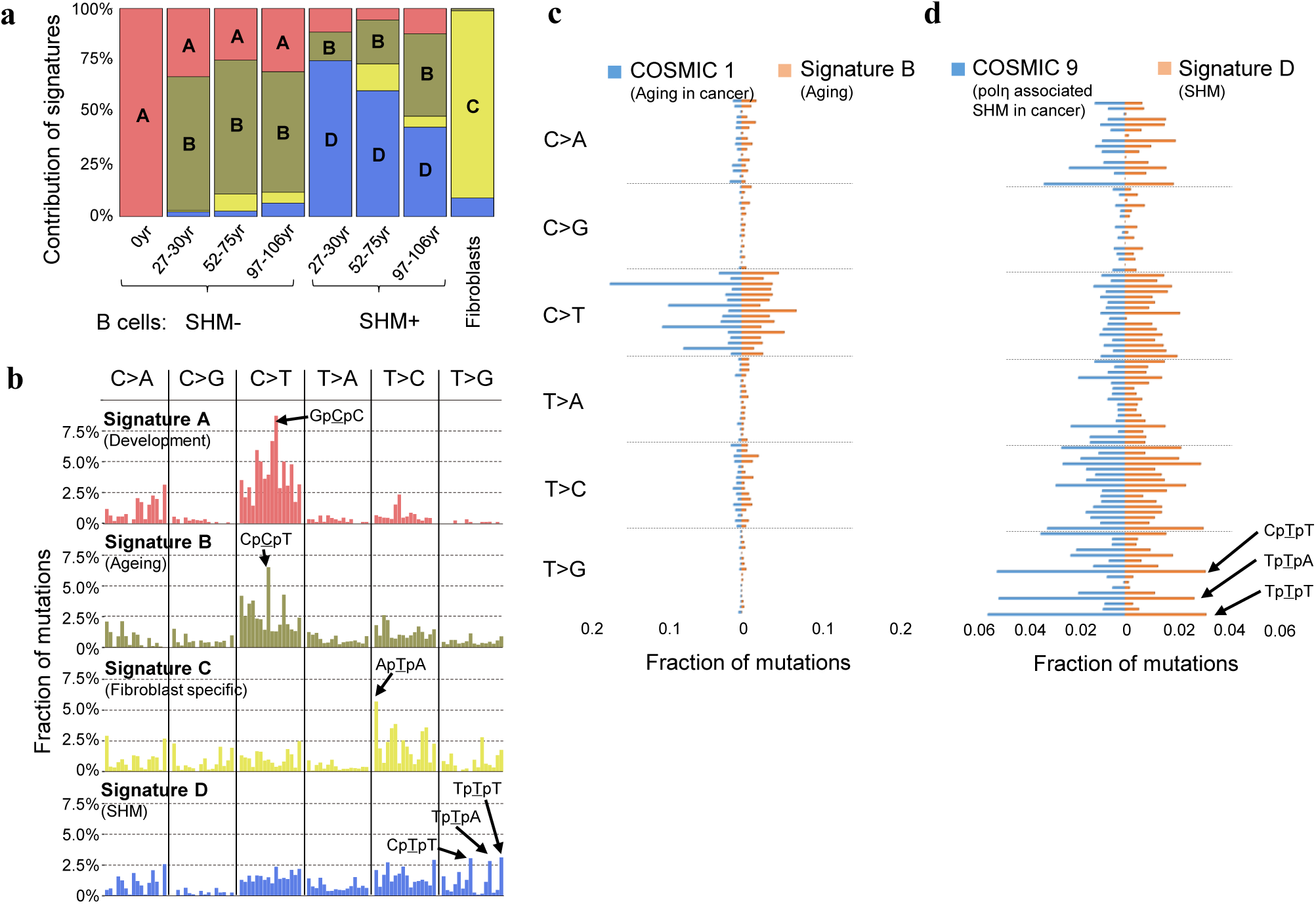
Mutation signatures. **a.** Contributions of signatures A, B and D to all mutations in B lymphocytes in the different age groups. Signature C is from spontaneous mutations previously obtained for human primary fibroblasts. **b.** Mutation signatures in the context of their flanking base pairs (in alphabetical order, e.g., in the first column C>A category from left to right, ApCpA > ApApA to TpCpT > TpApT). **c.** Comparison between signature B and COSMIC1. The fractions of mutations are presented in alphabetical row, e.g., in the first column C>A category from top down, ApCpA > ApApA to TpCpT > TpApT). **d.** Comparison between signature D and COSMIC 9.

Importantly, these signatures observed in normal B lymphocytes from healthy human donors were very similar to mutation signatures obtained from human cancers, i.e. Catalogue Of Somatic Mutations in Cancer (COSMIC; Supplementary Fig. 11d)^4,33,34^. Our ageing signature B was found to correlate highly with the cancer ageing signatures COSMIC1 and COSMIC5 in many cancer types (Spearman’s correlation coefficients ρ=0.80 and 0.74 separately, Fig. 2c, and Supplementary Fig. 11d). Our developmental signature A also correlates to COSMIC1 (ρ=0.78), as does signature B. These results suggest that the mechanistic origin of these signatures, even in tumors firmly resides in the normal process of age-related mutation accumulation.

The “ageing signature” appeared mostly a result of spontaneous cytosine deamination^35^. The latter was also observed in the mutation signature of a childhood cancer, pilocytic astrocytoma (COSMIC19), resulting in a high correlation, ρ=0.81, with our signature B (Supplementary Fig. 11d). We also observed a high correlation between the SHM signature D and the chronic lymphocytic leukemia and malignant B-cell lymphoma signature COSMIC9 (ρ=0.81, Fig. 2d). These results were confirmed by refitting the known COSMIC signatures to the observed somatic mutations in normal B lymphocytes, using the “deconstructSigs” package in R^36^. That is, COSMIC9 was only found in SHM+ cells and COSMIC5 and COSMIC19 were found in all cells (Supplementary Fig. 13). Together with the observation of off-target SHM hotspots, these results strongly support the hypothesis that somatic hypermutation contributes significantly to the development of B cell cancers^18^. No significant difference was found between the genome-wide mutation signatures and those in the 24 hotspot areas (Supplementary Fig. 14 in comparison with Fig. 2a and Supplementary Fig. 13).

An important question we were now able to directly address for the first time is whether somatic mutation accumulation during ageing can have a functional impact other than increasing cancer risk. For this purpose we subdivided all mutations into two groups: those that were and those that were not found in the functional part of the genome. Mutations in the functional genome were defined as those occurring in the transcribed exome or its regulatory regions. To identify the transcribed exome we performed RNA-seq of purified, bulk B lymphocytes (Supplementary Fig. 15, Supplementary Table 5; Methods). As gene regulatory regions we considered both promoters of active genes and open chromatin regions, e.g., transcription factor (TF) binding regions, identified by ATAC-seq in the same bulk B cells (Supplementary Fig. 16, Supplementary Table 5; Methods).

Of the 9.2±6.8 nonsynonymous mutations on average per cell in the transcribed B cell exome of the ≥ 97-year olds (Table 1; Supplementary Table 6), 6.5±5.5 (71%) were functionally deleterious according to SIFT and PROVEAN (Supplementary Table 7)^37,38^. Many more potentially functional mutations occurred in noncoding regions of the genome (Table 1, Supplementary Table 6). When analyzing the TF-binding regions reported by ENCODE^39^ in our B lymphocytes, the number of mutations in these regions were found to increase with age from 79.6 to 435.9 on average per cell (*P*=8.79×10^−6^). More specifically, when analyzing our own ATAC data, specific for B lymphocytes, we found an almost 5-fold age-related increase in the number of mutations in active open chromatin regions (likely to be TF-binding regions specific in B cells), from 5.4 to 24.5 per cell (*P*=0.0199). Even more mutations were found collectively in proximate promoter (from −1500 to +500bp of TSS), 5’ UTR and 3’ UTR regions, i.e., 56.9±26.9, 4.9±5.7 and 13.6±6.5, respectively, per cell in the oldest subjects. Hence, in the functional part of the B cell genome mutations were found to increase from 27.0±18.3 per cell in cells from newborns to 85.1±36.9 per cell in cells from >97 year olds (*P*=0.000878, exponential model non-linear least squares regression).

**Table 1.** Average number of functional SNVs per cell.

| Class<sup>1</sup> | Age |  |  |  |
| --- | --- | --- | --- | --- |
|  | 0 year | 27-30 years | 52-75 years | 97-106 years |
| Total in the genome | 463.4±179.0 | 1181.9±484.7 | 2101.7±2106.1 | 3127.0±1237.1 |
| Nonsynonymous | 2.2±4.1 | 1.7±2.5 | 5.4±5.3 | 9.2±6.8 |
| LOF SNV<sup>2</sup> | 0.4±1.1 | 0.0±0.0 | 0.6±1.7 | 0.5±1.4 |
| Damaging<sup>3</sup> | 1.3±2.8 | 1.7±2.5 | 4.0±5.0 | 6.5±5.5 |
| Open chromatin region<sup>4</sup> | 5.4±5.8 | 11.5±4.4 | 17.6±24.9 | 24.5±15.8 |
| Promoter | 17.7±10.1 | 19.6±7.4 | 33.1±34.9 | 56.9±26.9 |
| 5' UTR | 2.1±3.5 | 2.5±4.0 | 3.9±5.7 | 4.9±5.7 |
| 3' UTR | 4.6±5.9 | 11.0±11.9 | 10.9±11.2 | 13.6±6.5 |

<sup>1</sup>Except the genome total, only SNVs transcribed or in open chromatin regions are reported. A full table about both transcribed and non-transcribed SNVs is provided as Supplementary Table 6.
<sup>2</sup>Loss of function SNVs, including stop gain, stop loss and splicing alterations.
<sup>3</sup>SNVs annotated as damaging by SIFT or Deleterious by PROVEAN.
<sup>4</sup>Identified using ATAC-Seq of B lymphocytes.

Next, we tested if mutations in the functional genome were also functionally impactful. To do that we compared the pace of age-related accumulation of mutations in the functional genome with that of mutations in the genome overall. The results indicated a highly significant slower pace of potentially functional mutations (*P*=1.42×10^−14^, Fig. 3). In this case we used all protein-coding genes since we could not ascertain the origin of the mutations, which could have been in hematopoietic stem cells with other genes active than the mature B lymphocytes. These results indicate protection against deleterious mutations in the functional genome during human ageing, suggesting that many random somatic mutations are damaging to cellular function.

**Fig. 3.**
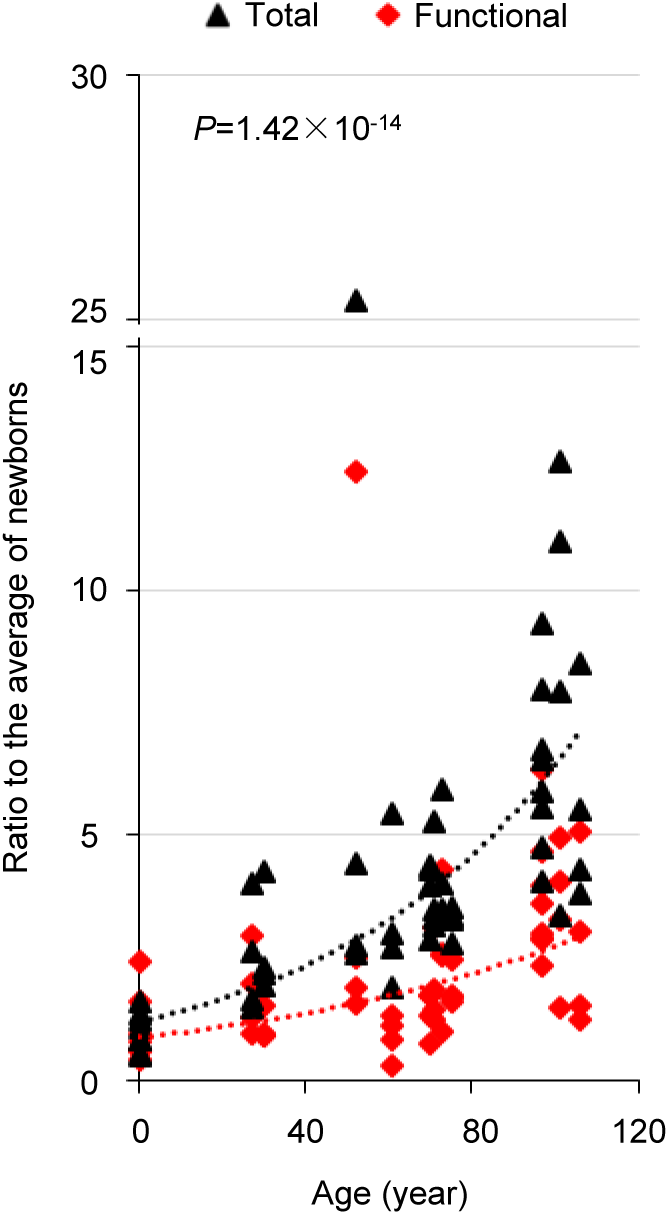
Accumulation of mutations in the functional genome and genome overall during ageing. Each data point represents the ratio of the number of mutations per cell to the average of the mutation number per cell in newborn B lymphocytes (functional genome and whole genome calculated separately). The ratios of the functional genome are in red and those in the genome overall in black. P value was estimated using Wilcoxon signed-rank test, two-tailed.

The significantly lower rate of mutation accumulation in functional genomic regions than the genome-wide average suggests that the B cells and their ancestral stem cells that harbored such mutations may have died or been at a growth disadvantage. This is in keeping with the discovery of a decreased number of lymphoid hematopoietic lineages during ageing^11^ and may underlie the CHIP phenomenon found in a fraction of elderly^40^. However, our data show that many mutations in the functional genome remain and continue to accumulate. While not lethal, such mutations may still contribute to the well-documented age-related decline in function that has been observed for many cell types in humans and experimental animals. As we and others have reported previously this effect may be mediated by increased cell-to-cell transcriptional variability^41-43^.

In summary, this first, comprehensive characterization of the landscape of somatic mutations in B lymphocytes across the entire human age range shows a highly significant age-related increase of mutations. Mutation signatures in these normal B cells, which were significantly different from those in human primary fibroblasts^3^, were found to correlate with those previously observed in human cancers, most notably B cell leukemia. This finding underscores the increasing realization that age is the most important risk factor for cancer. A substantial fraction of all mutations were found to occur in sequence features functionally active in B lymphocytes. These mutations were actively selected against as evidenced by their much slower rate of accumulation with age. Hence, somatic mutations may contribute to functional decline in ageing through depletion of cells or accumulation of cells with slightly deleterious mutations. Of note, in this study we only analyzed base substitution mutations. Other types of mutations such as small INDELS or genome structural variation are much less frequent, but may still contribute to a gradual age-related deterioration of the functional somatic genome. In principle we could test for such mutations in our single-cell whole-genome sequences. Reliable computational tools for that purpose, equally sensitive and accurate as our current tool for base substitutions are currently under development. Our present results provide the foundation for finally testing the somatic mutation theory of ageing by studying multiple human organs and tissues for the accumulation of potentially functional mutations and their causal relationship to age-related functional decline and disease.

## Acknowledgements

This study was supported by NIH grant P01 AG017242 (J.V.), K99 AG056656 (X.D.) and the Paul F. Glenn Center for the Biology of Human Aging. We thank the Molecular Cytogenetics Core and the Epigenomics Core at the Albert Einstein College of Medicine for B cell isolation and RNA-seq/ATAC-seq, respectively.

## Author Contributions

J.V., L.Z. and X.D. conceived this study and designed the experiments. L.Z. and M.L. performed the experiments. X.D. and T.W. analyzed the data. J.V., L.Z., X.D., T.W. and A.Y.M. wrote the manuscript.

