## Supplementary Methods for "Single-cell whole-genome sequencing reveals the functional landscape of somatic mutations in B lymphocytes across the human lifespan"

#### Human Specimens

Collection of blood for B lymphocyte isolation and whole-genome sequencing was approved by the Einstein-Montefiore institutional review board (IRB). Two frozen human umbilical CBMCs were obtained from AllCells Inc. (Alameda, CA) and StemCell Technologies Inc. (Vancouver, CA), separately. All donors in this study were healthy without a cancer history. Supplementary Table 1 shows age, gender and other information of the donors.

#### B lymphocyte isolation

For each donor, PBMCs were isolated using Ficoll-Paque PLUS (GE Healthcare) solution according to the manufacturer's instructions by the Molecular Cytogenetics Core at the Albert Einstein College of Medicine. Briefly, fresh peripheral blood was collected in ACD solution A and solution B anticoagulant tubes and diluted with PBS and 2 mM EDTA (ratio 1:1) (Gibco). Subsequently, the blood mixture was layered onto Ficoll-Paque PLUS. After centrifugation, lymphocyte cells remained at the plasma-Ficoll-Paque PLUS interface and were carefully transferred to a new tube and washed twice with 1x PBS. These PBMCs were re-suspended with MACS buffer (Miltenyi Biotec).

B lymphocytes were isolated using MACS separation (Kit #130-050301, Miltenyi Biotec). Briefly, each $10^7$ total PBMCs were re-suspended in 80 µl buffer and mixed with 20 µl CD19 MicroBeads. The mixture was incubated at 2-8 °C for 15 min, washed and centrifuged. Cell pellets were re-suspended and applied onto the column, which was placed in the magnetic field of a suitable MACS Separator. Unlabeled cells were washed away with washing buffer and collected for T cell isolation or discarded. Labeled B lymphocytes on the column were eluted into a collection tube.

To validate purity of B lymphocytes, cells were fluorescently stained with CD19-FITC (# 130-113-730, Miltenyi Biotec). Cell debris and dead cells were excluded from the analysis based on DAPI fluorescence. An example is shown in Supplementary Fig. 2a.

#### Single B lymphocyte isolation

Single B lymphocytes were collected using the CellRaft array (Cell Microsystems) following a protocol that was modified as compared to our previous procedure[1]. Briefly, a CellRaft array was rinsed with medium and subsequently coated with gelatin (2% in water; G1393-100ML Sigma-Aldrich) at 37°C for one hour to help the B lymphocytes attach[2,3]. After rinsing the superfluous gelatin solution off the array, approximately 5,000 B lymphocytes in 3 ml medium were added to the CellRaft array and incubated for 3 h at 37°C, 3% $O_2$, 10% $CO_2$. We washed the array twice with 1 ml of PBS to remove floating cells and added 3 ml of fresh complete medium. Single rafts containing one cell (as observed by microscopy using a 10x objective, (Supplementary Fig. 2b) were transferred, using the magnetic wand supplied with the CellRaft system, into 0.2 ml PCR tubes containing 2.5 µl PBS. We verified if there was only one raft in a PCR tube using a magnifying glass. Tubes containing single rafts were frozen immediately on crushed dry ice and kept at -80°C until use.

**Single-cell multiple displacement amplification**

We amplified four single B lymphocytes from each of 14 humans according to our SCMDA protocol[1]. Briefly, one µl Exo-Resistant random primer (ThermoFisher) was added to the cell, immediately followed by 3 µl lysis buffer (400 mM KOH, 100 mM DTT, 10 mM EDTA). Cell lysis and DNA denaturation were on ice for 10 min. Then 3 µl of stop buffer (400 mM HCl, 600 mM Tris-HCl, pH 7.5) was added to neutralize the lysis buffer. Finally, 32 µl of master mix containing 30 µl of MDA reaction buffer and 2 µl of Phi29 polymerase (REPLI-g UltraFast Mini Kit, Qiagen) was added. MDA was carried out for 1.5 h at 30°C; 3 min at 65°C, and kept at 4°C until purification. MDA product was purified using AMPureXP-beads (Beckman Coulter) and the concentration measured with the Qubit High Sensitivity dsDNA Kit (Invitrogen Life Science). Meanwhile, 1 ng of human genomic DNA in 2.5 µl PBS was amplified as a positive control, and 2.5 µl of PBS without any template was amplified as a negative control.

**Bulk DNA extraction**

Total DNA was collected from the cell pellets remaining after Ficoll density centrifugation for B lymphocyte isolation using the DNeasy Blood & Tissue Kit (Qiagen) following the manufacturer's specifications. The concentrations of DNA were quantified using the Qubit High Sensitivity dsDNA Kit (Invitrogen Life Science) and the qualities of DNAs were evaluated with 1% agarose gel electrophoresis.

**DNA library preparation and whole genome sequencing**

The libraries of 14 bulk DNAs and 56 single B lymphocyte MDA products were prepared using Truseq Nano DNA HT Sample Preparation Kit (Illumina USA) by Novogene (Beijing, China). Briefly, a total amount of 1.0 µg DNA per sample as input was fragmented by sonication to a size of 350 bp, and then DNA fragments were endpolished, A-tailed, and ligated with the full-length adapter for Illumina sequencing with further PCR amplification. PCR products were purified (AMPure XP system) and libraries were analyzed for size distribution by the Agilent 2100 Bioanalyzer and quantified using real-time PCR. All libraries were sequenced on the Illumina sequencing platform X Ten and paired-end reads, 2x150 bp were generated by Novogene, Inc.

**Alignment for whole-genome sequencing**

For each single cell or bulk sample, sequencing reads were adapter- and quality-trimmed using Trim Galore (version 0.3.7). Reads before and after trimming were subjected to quality checking using FastQC (version 0.11.4). The trimmed reads were aligned to the human reference genome (build 37) using BWA MEM (version 0.7.10)[4]. Sequence duplications were removed using samtools (version 0.1.19)[5]. The alignment was further indel-realigned based on known indels from the 1000 Genomes Project (phase I)[6], and their base quality scores were recalibrated based on known indels from the 1000 Genomes Project (phase I) and SNVs from dbSNP (build 144), both using GATK (version 3.5.0)[7].

**Calling somatic SNVs**

We used SCcaller (version 1.2) for identifying somatic mutations following its online instructions (https://github.com/biosinodx/SCcaller), requiring a minimum 20x sequencing depth and at least 4 reads containing the mutation[1]. As illustrated in the previous paper, the SCcaller identify somatic mutations from single cells by filtering out potential amplification artifacts. The filtering is based on a kernel smoother for estimating local allelic amplification bias and a likelihood ratio test to separate three models, one for heterozygous mutation, one for homozygous mutation and the other for the potential artifacts, at every locus. Because the SCcaller requires heterozygous SNPs to correct for allelic amplification bias, we only called mutations on autosomes. The heterozygous SNPs were identified in the bulk whole genome sequences using haplotypecaller of GATK.

**Estimating sensitivity of SNV calling**

The sensitivity of *de novo* mutation calling in the single cells was estimated as the ratio of the number of heterozygous SNPs to the total number of SNPs detected in bulk DNA, at a minimum sequencing depth of 20x in both single cells and bulk (Supplementary Table 3).

**Estimating specificity of SNV calling**

To validate variant calling, we performed Sanger sequencing on 20 randomly selected variants, 30 variants located at mutation clusters at immunoglobulin H chain and immunoglobulin L chain genes and 15 recurrent variants (identified in more than one cells) from the 56 single B lymphocytes.

**Detection of clonal hematopoiesis (CHIP) from bulk sequencing**

Besides looking at recurrent mutations from single cells, we also search from bulk sequencing for possible somatic CHIP mutations with variant allele frequencies significantly less than germline at 50%. First, candidate variants from bulk sequencing (the majority of which are germline) were called using GATK (version 3.5.0)[7], HaplotypeCaller (option: -stand_call_conf 30 -stand_emit_conf 10) without additional stringent filtering to include as many candidates as possible. Second, we annotated the candidate variants with ANNOVAR[8]. Third, using the ANNOVAR's annotations, we compared our variants with the CHIP variant provided by ref[9] Table S2, and kept any overlapping ones. Finally, we tested if any of the remaining variants has a variant allele frequency significantly differentiate from germline (50%) according to the method described in ref[9].

**Detection of mutational hotspots**

For detecting potential hotspot regions, we pooled the mutations identified in the 56 single B cells, in total 30,426 SNVs. We then applied the "shower" method using R package "ClusteredMutations"[10-12], requiring a minimum of 4 mutations within a distance that is less than or equal to 5kb. The average distance between two random, neighboring mutations is ~100kb. To calculate the false discovery rate (FDR) of the hotspots identified, we randomly distributed the 30,426 SNVs across the genome and applied the same criteria to discover potential hotspots to calculate the chance of false positives. This random process was repeated 2,000 times and resulted in the FDR estimations provided in Supplementary Table 4.

**Analyzing non-immunoglobulin mutation hotspots**

We annotated the 19 non-immunoglobulin hotspots through ENSEMBLE, ENCODE and by searching the literature. Eight of the 19 off-target hotspots (42%) were previously reported being off-target SHM hotspots in B cell-related cancers in humans or mice (Supplementary Table 4), with 6 of the 19 (32%) overlapping with specific TF binding sites, including E-Box binding regions, YY1 or C/EBP target regions associated with AID binding[13] (Supplementary Table 4). Using ATAC sequencing of B cells collected from two of our donors, we found the off-target hotspots significantly more likely to be in or close to (<1kb) open chromatin regions (3.8-fold more likely than random, $P$=0.003 with 2,000 permutations). We also found that significantly more off-target hotspots were close to transcription start sites (TSS; <1kb, 3.3-fold more likely than random, Monte Carlo $P$=0.013 with 2,000 permutations). A full annotation of all hotspots is provided as Supplementary Table 4.

**The "outlier" cell**

A SHM+ B cell from the 52-year old donor has a substantially higher mutation frequency than any other cell in this study (Fig. 1a). This cell has a stop codon loss of gene *HIST1H1E*, which is one of the most frequently mutated genes found in human lymphoma[14,15]. When performing the regression analysis in Supplementary Fig. 10a, we excluded this cell because it is the only SHM+ cell sequenced of the 52-year donor and is unlikely to be representative for mutation frequency of healthy SHM+ cells of this donor. However, this cell was included in all the other analyses in our study because in all the other analyses, this cell is not the only cell representing one donor or one group.

**Identifying CSR**

CSR can be identified as depletions of both IGHM and IGHD loci in the genome[16]. We first calculated the sequencing depths of the IGHM and IGHD loci as the average sequencing depths of all base pairs of the loci for every single B cell. We then calculated normalized depths as the ratio of the sequencing depths to genome-wide average sequencing depths of the same cells. To reliably distinguish CSR+ and CSR- cells, we applied an unsupervised method on the normalized depths as follows. We performed a principal component analysis (PCA) on the normalized depths of the two loci. All cells clearly separated into two groups based on values of the first principle component (PC1; Supplementary Fig. 5a). We then modeled the distribution of PC1 values as a mixture of two normal distributions estimated using an Expectation–Maximization algorithm from R package "mixtools" (Supplementary Fig. 5b)[17]. Posterior probabilities of every single cell as a member of either of the two normal distributions were calculated using the same R package (Supplementary Fig. 5c). Based on the posterior probabilities, we distinguished CSR+ and CSR- cells as indicated in Supplementary Fig. 5c.

**Analysis of mutation spectra of the affected bases**

The mutation spectra of human B lymphocytes were found to be dominated by mutations at cytosines and guanines and to differ significantly from those of the dermal fibroblasts analyzed previously (Supplementary Fig. 7a)[1]. More GC>AT transitions were observed in the B lymphocytes than in the fibroblasts (21%), with GC>AT transitions overall significantly more dominant in B lymphocytes from the newborns (66%) than in those from the other age groups (44% to 47%; $P$=2.18x$10^{-9}$, t-test, two-tailed).

Since the GC>AT transition is the major fraction of the spectra, we tested the correlation of CpG methylation and mutation frequency because cytosine in CpG dinucleotides is prone to deamination and methylated CpG is even more vulnerable without the protection of uracil glycosylase [18]. CpG methylation data at single cytosine resolution as determined by whole-genome bisulfite sequencing of adult B lymphocytes (CD19+) was obtained from the ENCODE (Supplementary Table 5; see the following paragraph). We found a significantly higher number of mutations at methylated CpGs than what would be expected by chance alone (Supplementary Fig. 7b). These results are similar to what has been found for germline mutations and somatic mutations in human tumors[19,20].

To analyze mutations in relation to methylated cytosines, we used data from ENCODE. Raw sequencing reads of whole-genome bisulfite sequencing of B cells (CD19+) of a 37yr old male donor were downloaded from ENCODE database (https://www.encodeproject.org/experiments/ENCSR284TCU) and were adapter and quality-trimmed using Trim Galore (version 0.3.7). Reads before and after trimming were subjected to quality checking using FastQC (version 0.11.4). First and second end of trimmed reads were aligned to the human reference genome (GRCh37.73) using Bismark (version 0.14.4) with the alignment tool Bowtie2 (version 2.2.3) separately[21,22]. Sequence duplicates were removed, and single CpG methylation called using Bismark, then CpG calls from the two ends of reads were merged with inconstant CpG calls of the two ends discarded. Single CpGs with at least 5 reads were considered for the methylation analysis in Supplementary Fig. 7b. Bisulfite conversion rate of the ENCODE experiment was estimated to be over 99%.

**Identifying mutation signatures**

We pooled mutations from the following groups of cells: cord blood, SHM- B cells of the 27-30yr old, SHM- B cells of the 52-75yr old, SHM- B cells of the 97-106yr old, SHM+ B cells of the 27-30yr old, SHM+ B cells of the 52-75yr old, SHM+ B cells of the 97-106yr old, and the 6 fibroblasts analyzed previously[1]. Using NMF decomposition in the R package "SomaticSignatures"[23], we identified 4 signatures from the above 8 groups. Thirty cancer mutation signatures from COSMIC were downloaded from http://cancer.sanger.ac.uk/cosmic/signatures and we calculated the Spearman's correlation coefficients ρ between COSMIC signatures and ours using R.

**RNA sequencing**

We performed RNA sequencing on frozen B lymphocytes from two donors (F1 and M2; Supplementary Table 1), in duplicate. For each donor, the two replicates were from two blood draws on separate days. Total RNA of B lymphocytes was extracted using RNeasy Micro Kit (Qiagen) according to the manufacturer's specification. The concentrations of RNA were quantified with Qubit RNA HS Assay Kit (Invitrogen Life Science) and the qualities of RNA were evaluated using bioanalyzer with Agilent RNA 6000 Pico Kit (Agilent Technologies). The RIN number of each sample submitted for sequencing was about 9.0. Libraries were prepared from 500 ng RNA using KAPA Stranded RNA-Seq Kit with RiboErase (KK8483, KAPA Biosystems) with Trueseq Index (Illumina) by the Einstein Epigenomics Core. Briefly, rRNA was depleted by hybridization with complementary DNA oligonucleotides, treated with RNase H and DNase to remove rRNA duplexed to DNA and the DNA oligonucleotides. RNA was fragmented using heat and magnesium. Subsequently, $1^{st}$ strand and $2^{nd}$ strand were synthesized to generate double-stranded cDNA (dscDNA). A-tailing was added to 3'-ends of the dscDNA and ligated with adapters. The ligated library was amplified by PCR. The libraries of the four RNA samples were sequenced on the Illumina HiSeq 2500, with 2×100 bp paired-end reads.

After quality check using FastQC (version 0.11.4), raw sequence reads for each sample were aligned to human reference genome (GRCh37.73) using STAR (version 2.5.2b; options: --outSAMattrIHstart 0 --outFilterIntronMotifs RemoveNoncanonical --alignIntronMin 20 --alignIntronMax 1000000 --outFilterMultimapNmax 1 --outSAMtype BAM SortedByCoordinate)[24], and sequence duplications were filtered out using Picard tools (version 1.119). FPKM values of gene expression were calculated using cufflinks (version 2.2.1) with gene annotation from ENSEMBL database (GRCh37.73)[25]. Finally, genes with FPKM consistently larger than 1 in all four samples are considered as transcribed, and the other genes untranscribed.

**ATAC sequencing**

ATAC sequencing was performed on fresh B lymphocytes from F1 and M2 donors. Libraries were prepared as described[26]. Briefly, to isolate nuclei, we spun 25,000 B cells and washed once with cold PBS. Cells were lysed using 10 mM Tris-HCl, pH 7.4, 10 mM NaCl, 3 mM MgCl2, 0.1% IGEPAL CA-630[27]. The nuclei pellet was re-suspended in transposase reaction mix, including 25 µl of 2xTD buffer and 2 µl of Tn5 transposase (Nextera DNA Library Prep Kit, Illumina) and incubated for 30 min at 37°C. After transposition, the fragmented sample was purified using the MinElute PCR Purification Kit (Qiagen) and subsequently PCR-amplified, (10 ul of transposed DNA, 5 µl of nuclease-free water, 5 µl of primer N70#, 5 µl of primer N50#, 25 µl of NEB High Fidelity 2 x PCR Master Mix). PCR conditions were 72 °C for 5 min; 98 °C for 30 s; and five cycles 98 °C for 10 s, 63 °C for 30 s and 72 °C for 1 min. To reduce GC and size bias in PCR, the PCR reaction was monitored using qPCR with 4 µl of the PCR-amplified library, 1 µl of PPC (Nextera DNA Library Prep Kit, Illumina) and 5 µl of 2×SyBr Green Master (Applied Biosystems). The library was amplified for a total of 18 cycles. The libraries were purified using QIAquick PCR Purification Kit (Qiagen) and quantified using Qubit High Sensitivity dsDNA Kit (Invitrogen Life Science). Library quality was assessed using the Agilent Bioanalyzer High-Sensitivity DNA kit (Agilent Technologies). Libraries were sequenced on the Illumina HiSeq 2500, with 2×100 bp reads by the Einstein Epigenomics Core.

The raw reads of ATAC sequencing were adapter- and quality-trimmed using Trim Galore (version 0.3.7). Reads before and after trimming were subjected to quality checking using FastQC (version 0.11.4). Alignment to human reference genome (GRCh37.73) was performed using bowtie2 (version 2.2.3; option: -X 2000). Duplications were removed using Picard tools (version 1.119). Reads with low mapping quality (MapQ<30) were discarded. Open chromatin regions as ATAC sequencing peaks were called using MACS2 (version 2.1.1; option: callpeak -g hs --nomodel --shift -100 --extsize 200)[28].

Mapabilities of all the peaks were estimated using Mapability score obtained from the USCS genome browser and >95% of peaks of both samples were in regions with highest mapability (Mapability score=100%). We used bedtools to determine if reads overlap with the peaks (requiring >= 1 bp overlap), and then checked if the peaks overlap with known TF binding regions reported in ENCODE (requiring 50% bp of peaks overlap with TF binding region;

Supplementary Table 5). To check the consistency between our ATAC sequencing and RNA sequencing results, we compared enrichment of ATAC seq peaks at TSS of transcribed and untranscribed genes (FPKM>=1 and FPKM<1 respectively, Supplementary Fig. 10b). The peak enrichment at TSS was calculated using HOMER (http://homer.ucsd.edu/homer/).

Finally, we merged the open chromatin peaks called from sample F1 and M2 using bedtools (version 2.25.0)[29] to annotate somatic mutations.

### Data availability

The sequencing data is being uploaded to SRA database.

1
