## Supplementary Figures for "Single-cell whole-genome sequencing reveals the functional landscape of somatic mutations in B lymphocytes across the human lifespan"

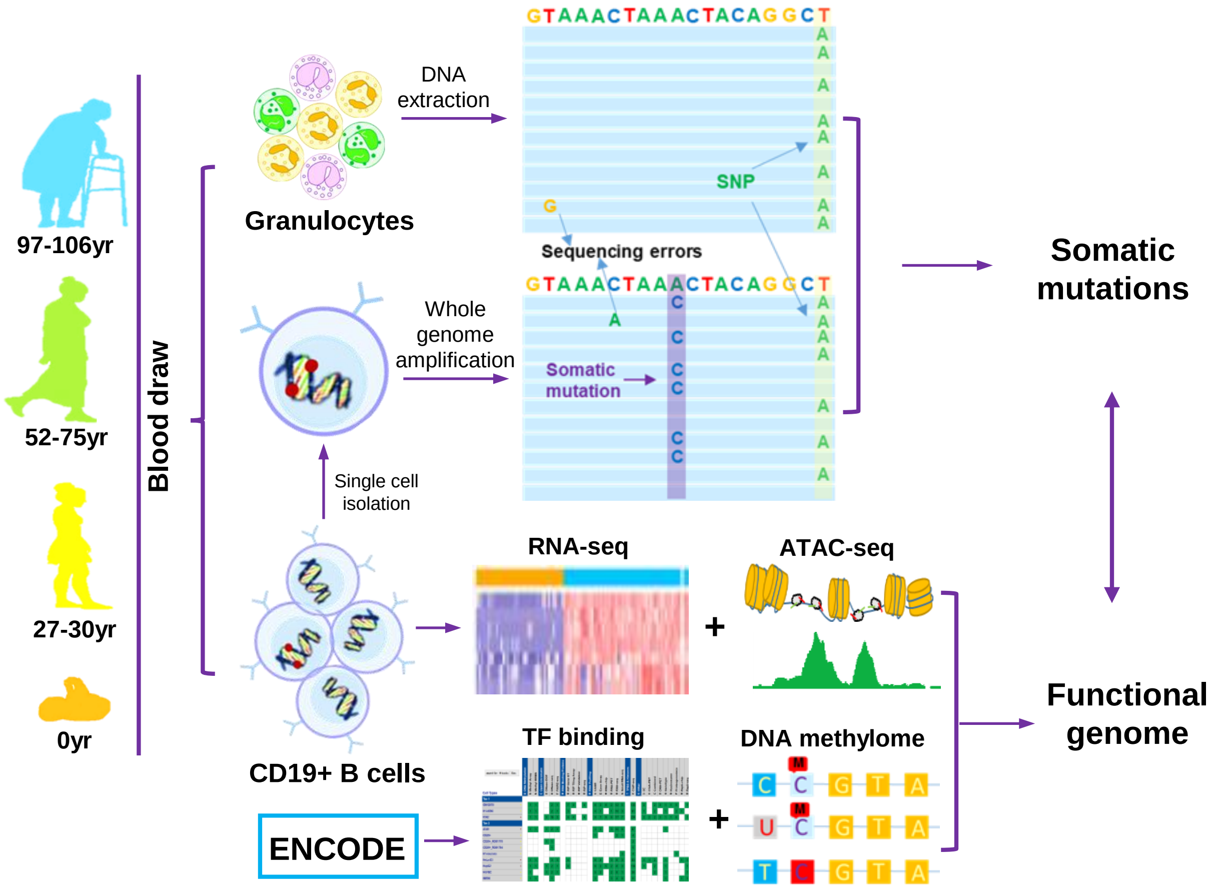


**Supplementary Fig. 1 | Study design.**
A schematic illustration of the single-cell approach to characterize the somatic mutational landscape as a function of age in human B lymphocytes.

**a**

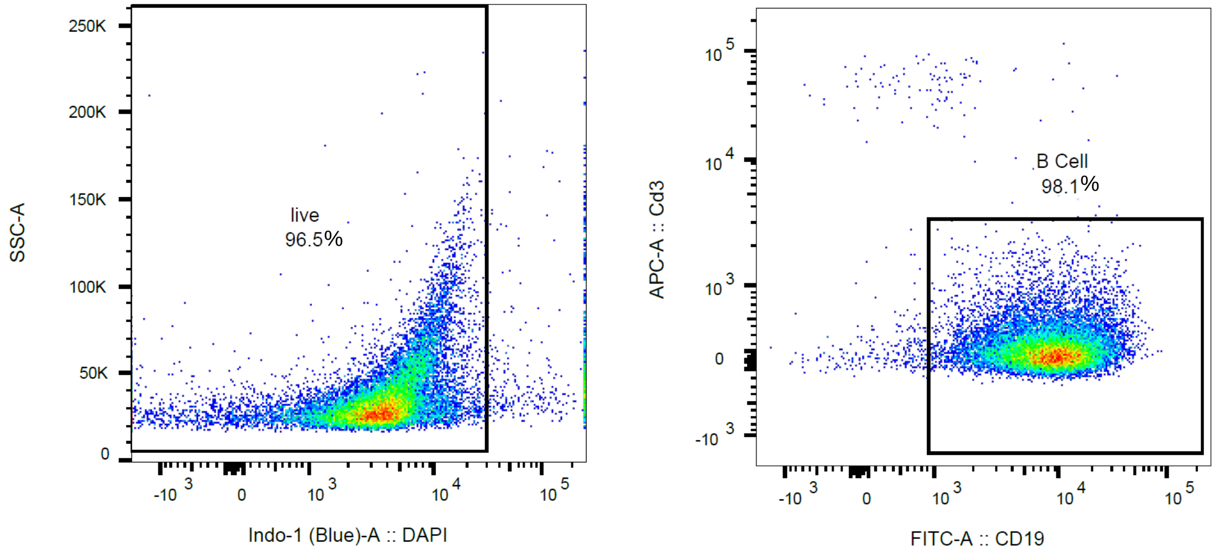


**b**

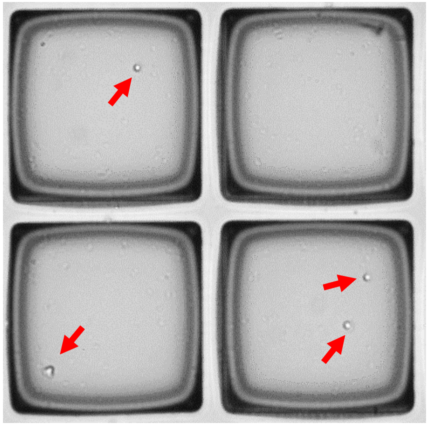

**Supplementary Fig. 2 | Isolating single B lymphocytes.**
**a.** Specificity of B cells isolated using MACS was validated using FACS with DAPI staining for viability checking as well as APC CD3 and FITC CD19 staining for B cell checking. **b.** Isolating single B cells using CellRaft. The red arrows indicate single cells.

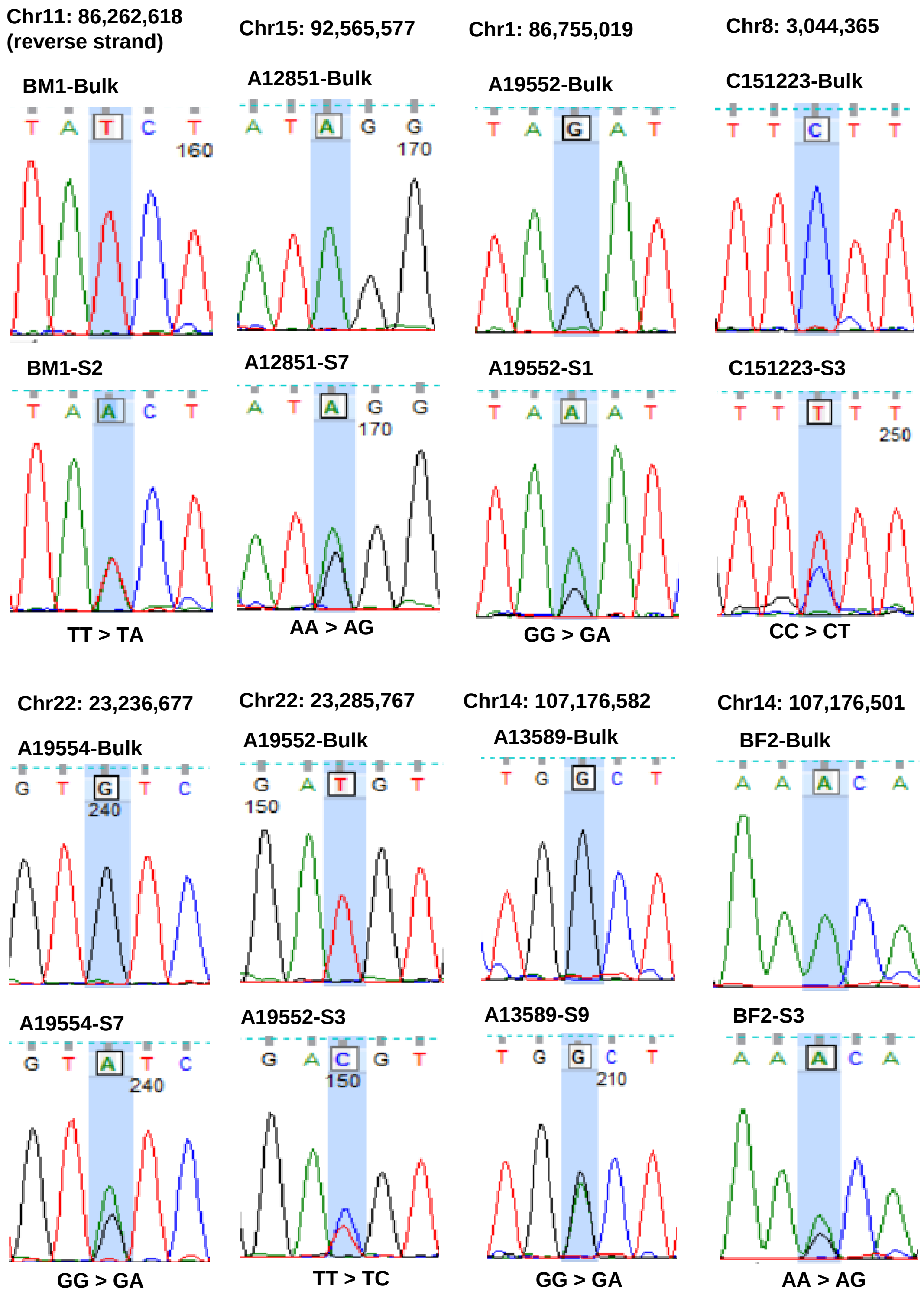


**Supplementary Fig. 3 | Sanger sequencing of randomly selected SNVs.**
The genome coordinates of mutations are provided at the top, followed by results of bulk, and results of the single cell.

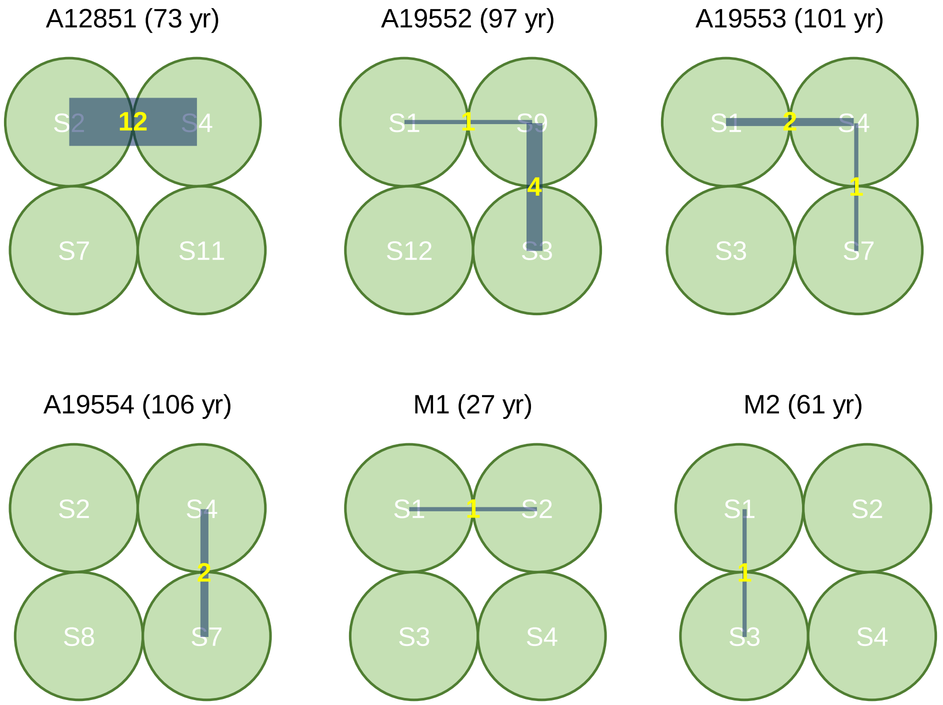


**Supplementary Fig. 4 | Recurrent mutations.**
A circle presents a single cell. A line between two cells with a number indicates the number of recurrent mutations found in the two cells.

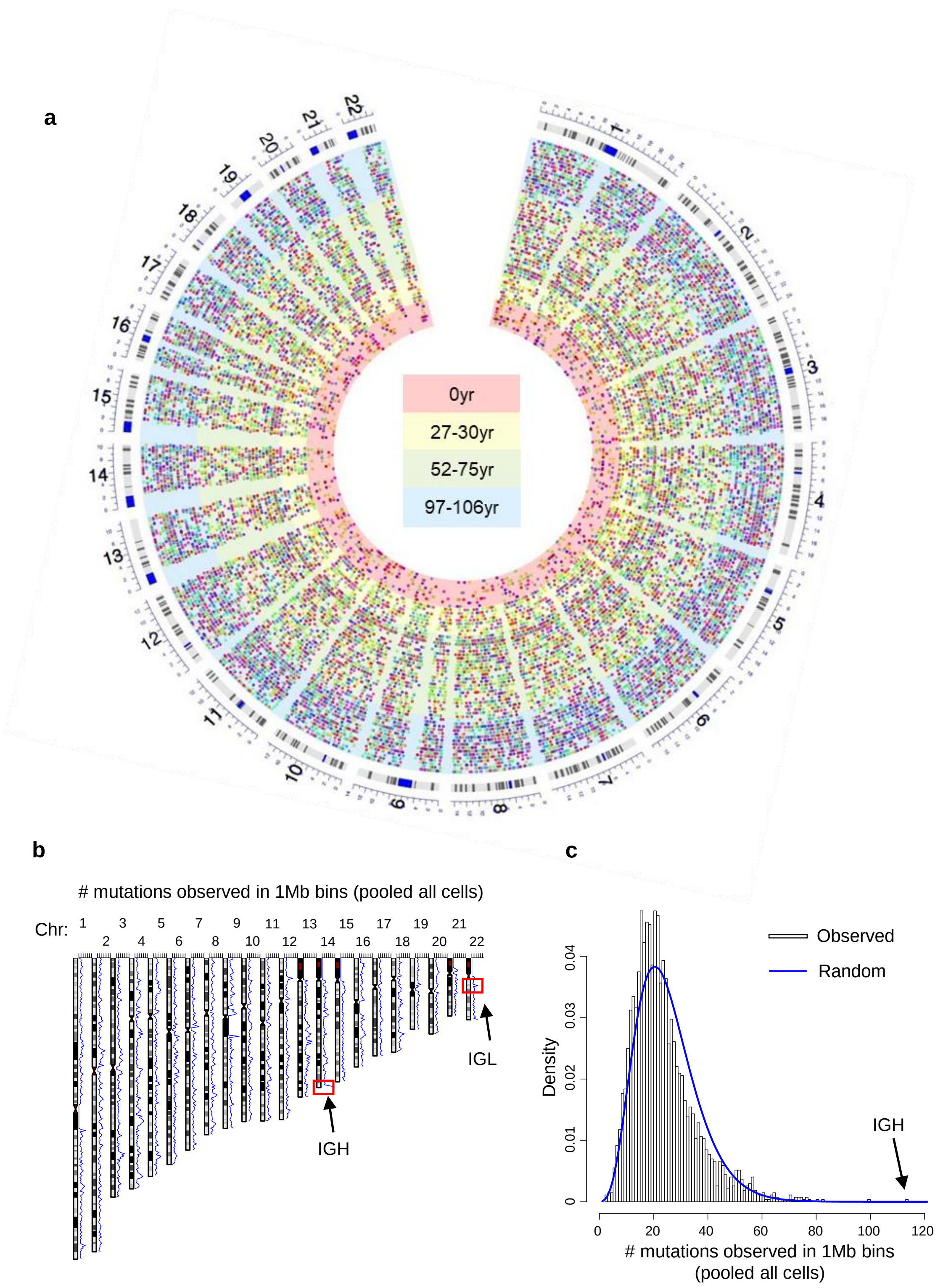


**Supplementary Fig. 5 | Distribution of somatic mutations at Mb scale.**
**a.** Circus diagram illustrates genome-wide distribution of SNVs. Each circle represents one cell. Dots present SNVs. Neighboring SNVs are plotted with different colors. Panels correspond to age groups of donors. **b.** Number of mutations (the blue lines) observed in Mb bins across the genome. Mutations from all 56 B cells were pooled in this analysis. **c.** Distribution of the numbers of mutation per Mb bin. The blue line represents the expected negative binomial distribution, which describes random distribution.

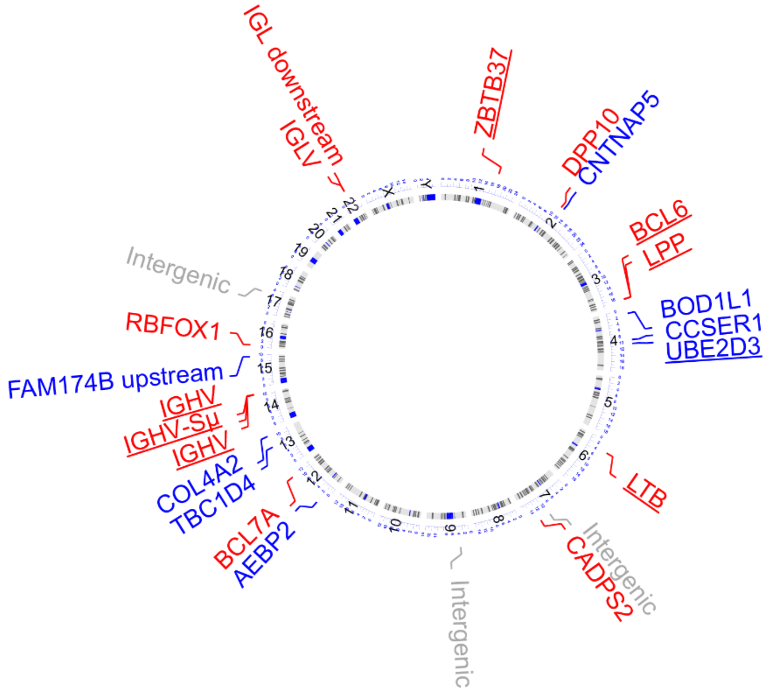


**Supplementary Fig. 6 | Distribution of mutation hotspots across the genome.** Hotspot names in red and blue indicate the hotspots that previously reported in human studies and those that have been reported separately (Supplementary Table 4 for details). Hotspot names underlined indicates the hotspots that were discovered in mouse an AID ChIP-sequencing study (see Supplementary Fig. 8).

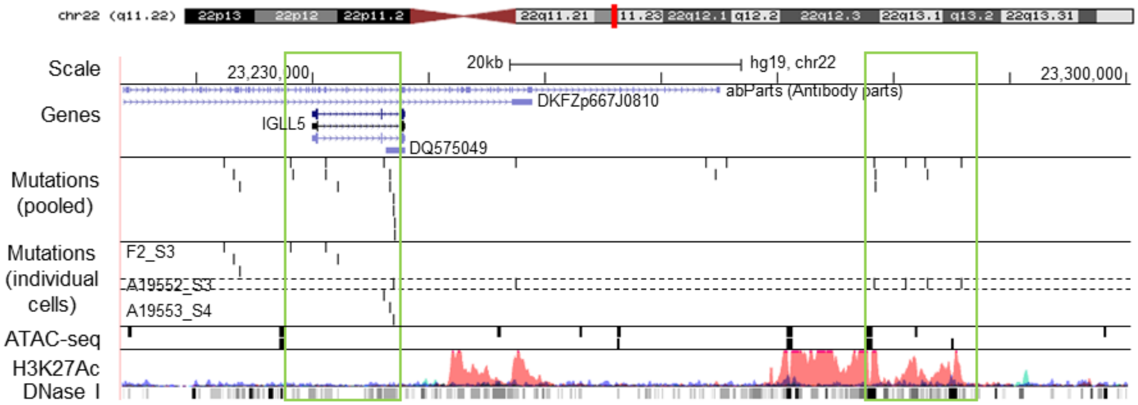


**Supplementary Fig. 7 | Mutation hotspots in IGL regions visualized using the UCSC genome browser.**
The legend is the same as Fig. 1c.

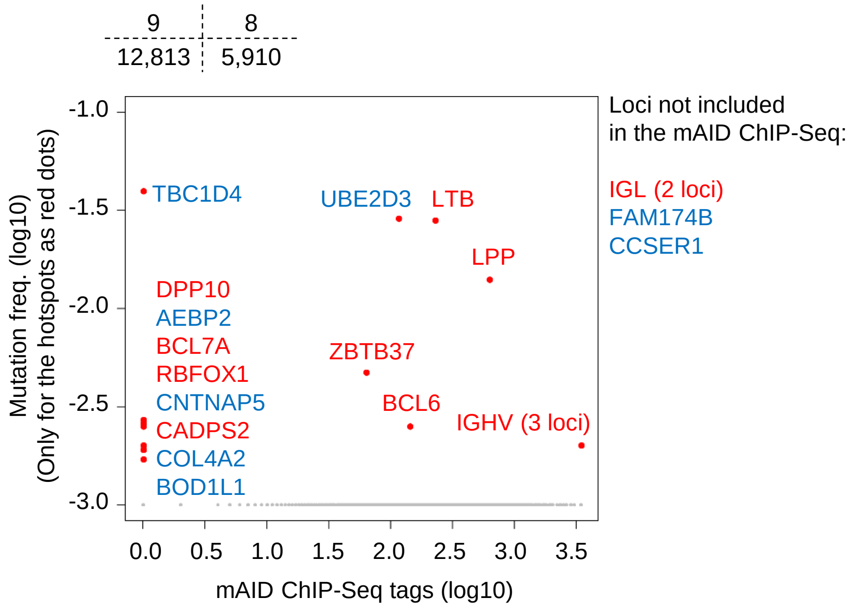


**Supplementary Fig. 8 | Comparing the hotspots with mAID ChIP-sequencing data.** The red dots indicate hotspots identified in this study. The blue dots indicates all the other genes with the mouse AID ChIP-sequencing data. The red and blue colour code of hotspot name is the same as Supplementary Fig. 6.

**a**

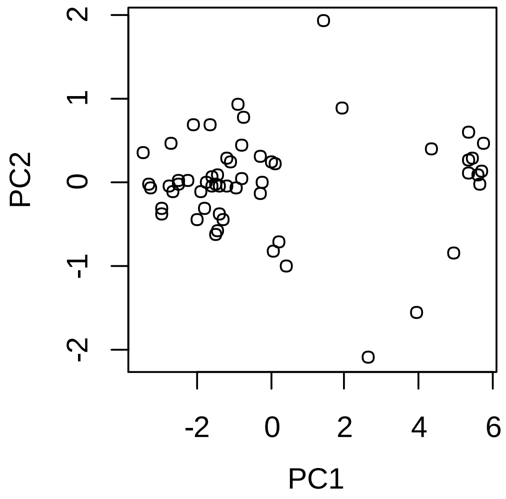


**b**

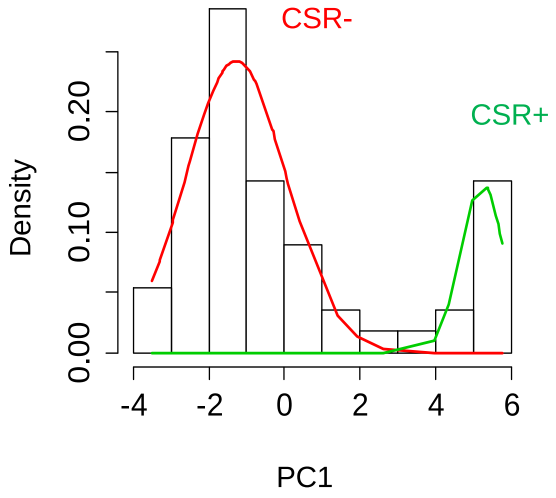


**c**

| SHM+ cells | CSR+ cells |
|---|---|

| Cell ID | Prob. of CSR- | Prob. of CSR+ |
|---|---|---|
| C151223_S1 | 1.00E+00 | 1.21E-26 |
| C151223_S2 | 1.00E+00 | 2.63E-30 |
| C151223_S3 | 1.00E+00 | 1.90E-19 |
| C151223_S4 | 9.99E-01 | 6.33E-04 |
| C70007_S1 | 1.00E+00 | 2.90E-17 |
| C70007_S10 | 1.00E+00 | 2.38E-33 |
| C70007_S3 | 1.00E+00 | 1.28E-34 |
| C70007_S5 | 1.00E+00 | 5.52E-36 |
| BM1_S1 | 4.50E-06 | 1.00E+00 |
| BM1_S2 | 1.00E+00 | 1.60E-07 |
| BM1_S3 | 1.00E+00 | 1.33E-29 |
| BM1_S4 | 1.00E+00 | 1.57E-43 |
| BF1_S1 | 2.89E-06 | 1.00E+00 |
| BF1_S2 | 1.00E+00 | 2.85E-38 |
| BF1_S3 | 1.00E+00 | 7.88E-28 |
| BF1_S4 | 1.00E+00 | 4.33E-34 |
| BF2_S1 | 1.00E+00 | 2.00E-32 |
| BF2_S2 | 1.00E+00 | 2.34E-34 |
| BF2_S3 | 5.79E-06 | 1.00E+00 |
| BF2_S4 | 1.00E+00 | 2.87E-46 |
| BM2_S1 | 1.00E+00 | 5.57E-52 |
| BM2_S2 | 1.00E+00 | 5.85E-33 |
| BM2_S3 | 1.00E+00 | 1.82E-30 |
| BM2_S4 | 1.00E+00 | 1.65E-32 |
| A19505_S1 | 1.00E+00 | 5.37E-32 |
| A19505_S2 | 1.00E+00 | 3.02E-43 |
| A19505_S7 | 1.00E+00 | 1.90E-10 |
| A19505_S8 | 1.00E+00 | 2.64E-48 |

| Cell ID | Prob. of CSR- | Prob. of CSR+ |
|---|---|---|
| A13589_S10 | 1.00E+00 | 1.60E-45 |
| A13589_S3 | 1.00E+00 | 4.30E-48 |
| A13589_S6 | 1.00E+00 | 3.70E-52 |
| A13589_S9 | 1.00E+00 | 3.29E-39 |
| A12851_S11 | 1.00E+00 | 1.59E-32 |
| A12851_S2 | 1.00E+00 | 4.26E-22 |
| A12851_S4 | 1.00E+00 | 2.68E-31 |
| A12851_S7 | 9.84E-03 | 9.90E-01 |
| A11365_S1 | 1.00E+00 | 1.41E-22 |
| A11365_S2 | 1.00E+00 | 3.90E-28 |
| A11365_S3 | 1.00E+00 | 6.41E-37 |
| A11365_S4 | 1.00E+00 | 2.10E-35 |
| A19552_S1 | 1.00E+00 | 2.21E-26 |
| A19552_S12 | 1.00E+00 | 1.85E-18 |
| A19552_S3 | 1.91E-06 | 1.00E+00 |
| A19552_S9 | 1.00E+00 | 1.16E-22 |
| A20061_S14 | 1.00E+00 | 5.93E-41 |
| A20061_S16 | 2.50E-06 | 1.00E+00 |
| A20061_S2 | 1.00E+00 | 1.93E-54 |
| A20061_S9 | 1.00E+00 | 7.79E-45 |
| A19553_S1 | 1.00E+00 | 8.45E-27 |
| A19553_S3 | 2.66E-05 | 1.00E+00 |
| A19553_S4 | 5.66E-06 | 1.00E+00 |
| A19553_S7 | 2.18E-06 | 1.00E+00 |
| A19554_S2 | 6.80E-04 | 9.99E-01 |
| A19554_S4 | 1.00E+00 | 9.14E-20 |
| A19554_S7 | 5.43E-06 | 1.00E+00 |
| A19554_S8 | 1.00E+00 | 3.58E-20 |

**Supplementary Fig. 9 | Identifying CSR from single cells.**
**a.** PCA of normalized depths of IGHM and IGHD loci. **b.** Distribution and modeling of PC1 of the PCA. It virtually separates CSR+/- cells. **c.** Posterior probability of being CSR+/- cells.

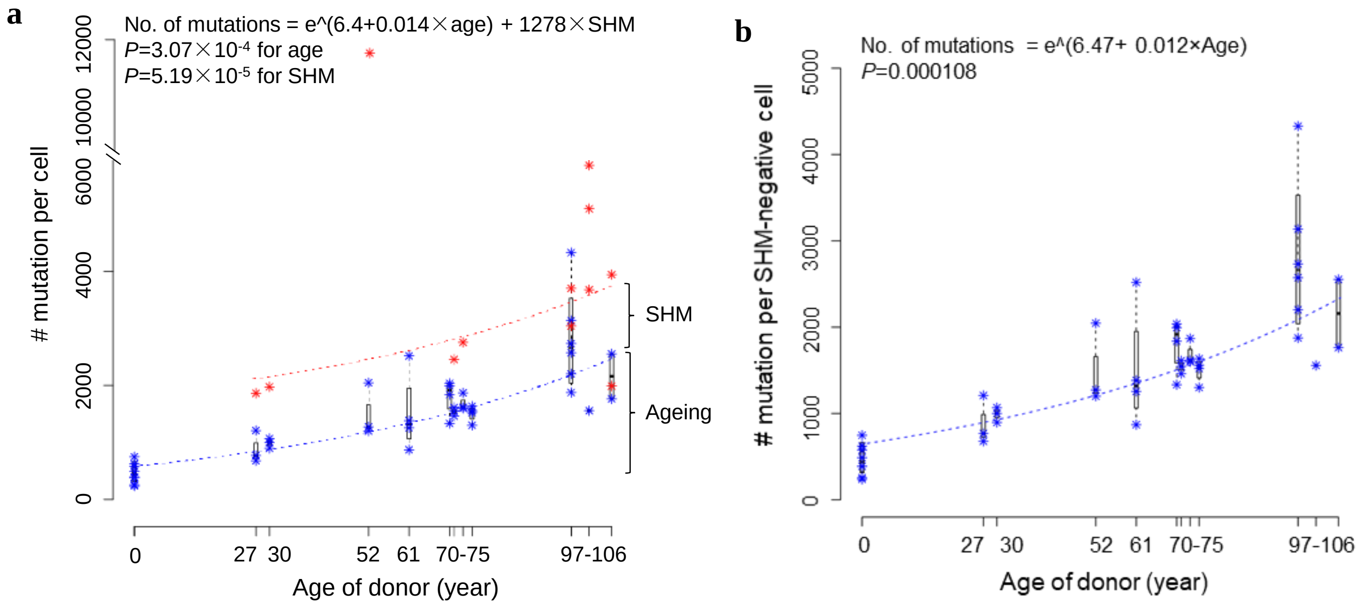


**Supplementary Fig. 10 | Mutation frequencies of SHM+/- B cells.**
**a.** Mutation frequency with age. SHM+ (red asterisks) or SHM- (blue asterisks) cells are considered separately. Regression analyses were performed on the median mutation frequency of all cells of donors. The cell with the highest mutation frequency (from the 52-year old) was not included in the regression analysis (see supplementary text for more discussions about the "outlier" cell). **b.** Somatic SNV frequency in SHM- cells. Both regressions in **a** and **b** were performed using R with the function "nls".

**a**

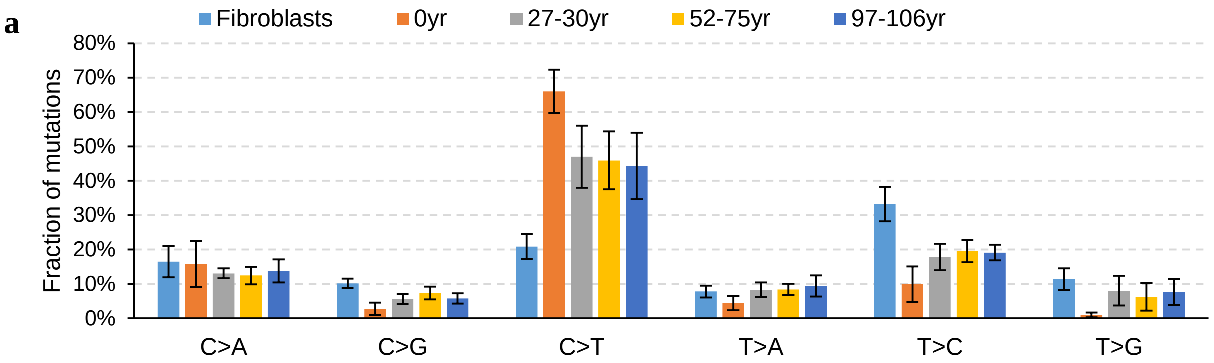


**b**

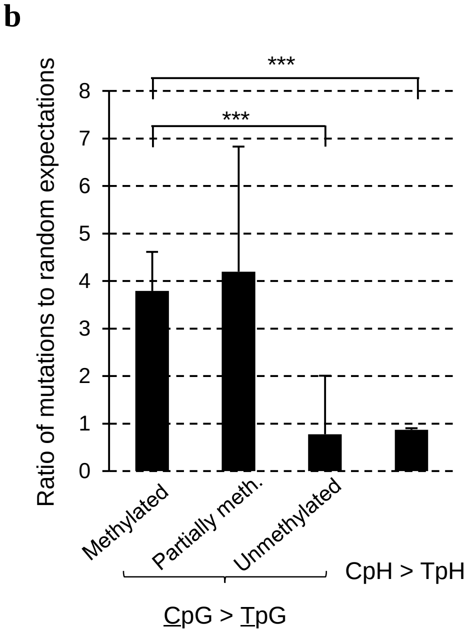


**c**

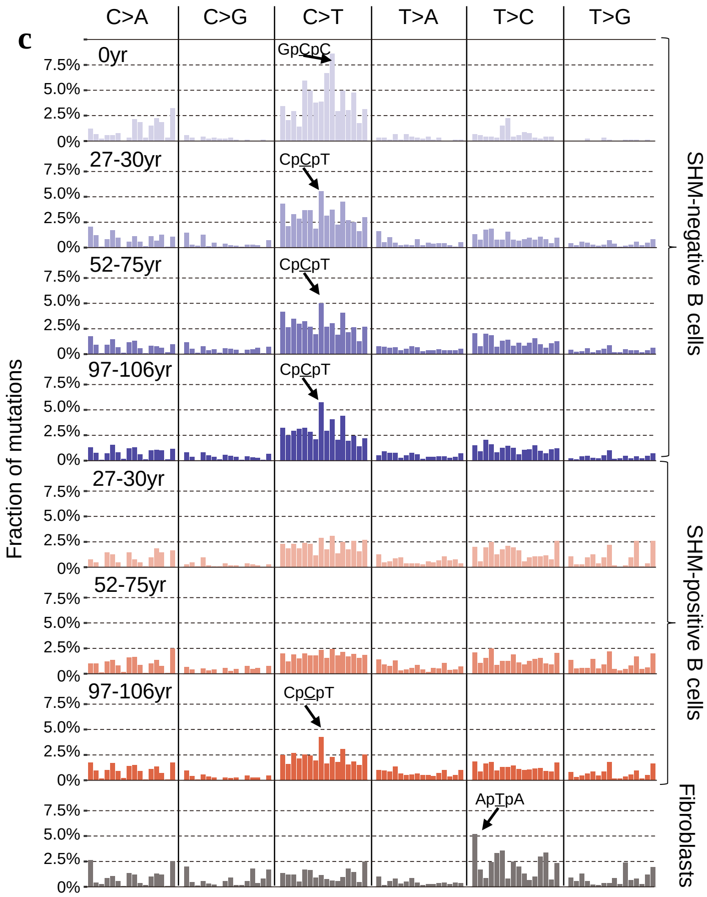


**d**

| | Sig. A | Sig. B | Sig. C | Sig. D |
|---|---|---|---|---|
| COSMIC1 | 0.78 | **0.80** | 0.43 | 0.61 |
| COSMIC2 | 0.62 | 0.60 | 0.39 | 0.28 |
| COSMIC3 | 0.25 | 0.36 | 0.42 | 0.11 |
| COSMIC4 | 0.45 | 0.40 | -0.04 | -0.11 |
| COSMIC5 | 0.69 | 0.74 | 0.52 | 0.63 |
| COSMIC6 | 0.68 | 0.65 | 0.21 | 0.52 |
| COSMIC7 | 0.33 | 0.42 | 0.32 | 0.46 |
| COSMIC8 | 0.38 | 0.47 | 0.31 | 0.43 |
| COSMIC9 | 0.22 | 0.20 | 0.51 | **0.81** |
| COSMIC10 | 0.42 | 0.36 | 0.32 | 0.60 |
| COSMIC11 | 0.68 | 0.71 | 0.23 | 0.33 |
| COSMIC12 | 0.53 | 0.56 | 0.45 | 0.57 |
| COSMIC13 | 0.30 | 0.33 | 0.12 | 0.06 |
| COSMIC14 | 0.69 | 0.64 | 0.28 | 0.46 |
| COSMIC15 | 0.73 | 0.66 | 0.33 | 0.49 |
| COSMIC16 | 0.44 | 0.50 | 0.56 | 0.60 |
| COSMIC17 | 0.30 | 0.31 | 0.32 | 0.36 |
| COSMIC18 | 0.51 | 0.46 | 0.10 | 0.12 |
| COSMIC19 | 0.75 | **0.81** | 0.37 | 0.51 |
| COSMIC20 | 0.68 | 0.64 | 0.25 | 0.44 |
| COSMIC21 | 0.48 | 0.56 | 0.30 | 0.46 |
| COSMIC22 | 0.30 | 0.32 | 0.05 | 0.33 |
| COSMIC23 | 0.64 | 0.68 | 0.18 | 0.26 |
| COSMIC24 | 0.40 | 0.34 | 0.03 | -0.16 |
| COSMIC25 | 0.37 | 0.45 | 0.38 | 0.46 |
| COSMIC26 | 0.51 | 0.56 | 0.41 | 0.54 |
| COSMIC27 | 0.68 | 0.59 | 0.32 | 0.38 |
| COSMIC28 | 0.07 | 0.06 | 0.31 | 0.46 |
| COSMIC29 | 0.58 | 0.50 | 0.26 | 0.26 |
| COSMIC30 | 0.39 | 0.38 | -0.11 | 0.18 |

**Supplementary Fig. 11 | Mutation spectra.**
**a.** Mutation spectra of B cells and fibroblasts. Each bar represents an average of all single cells of the group. Error bars represent s.d., **b.** Fractions of GC>AT transitions at CpG sites. Methylated, partial and unmethylated CpG sites were identified using whole-genome bisulfite sequencing data from ENCODE (Methods). Error bars indicate s.d. Both P values are smaller than $10^{-20}$, using paired t test two tailed. **c.** Mutation spectra in context of their flanking base pairs. SHM+ and SHM- cells were plotted separately. The x-axis labels are in alphabetical order, i.e., in the first column C>A category from left to right, ApCpA > ApApA to TpCpT > TpApT. **d.** Spearman's correlation coefficients between our signatures A to D and COSMIC1 to 30.

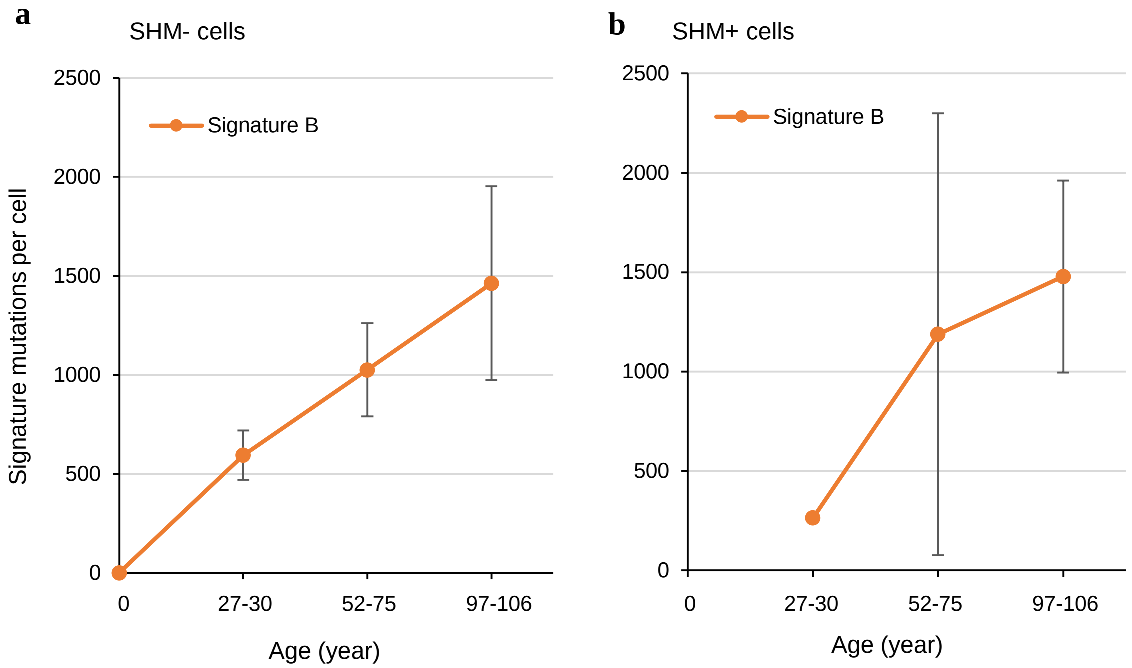


*P*=0.000925 for signature B
Linear region: No. signature mutation ~ age + SHM status

**Supplementary Fig. 12 | Numbers of signature mutations during ageing.**
**a.** SHM- cells. **b.** SHM+ cells.

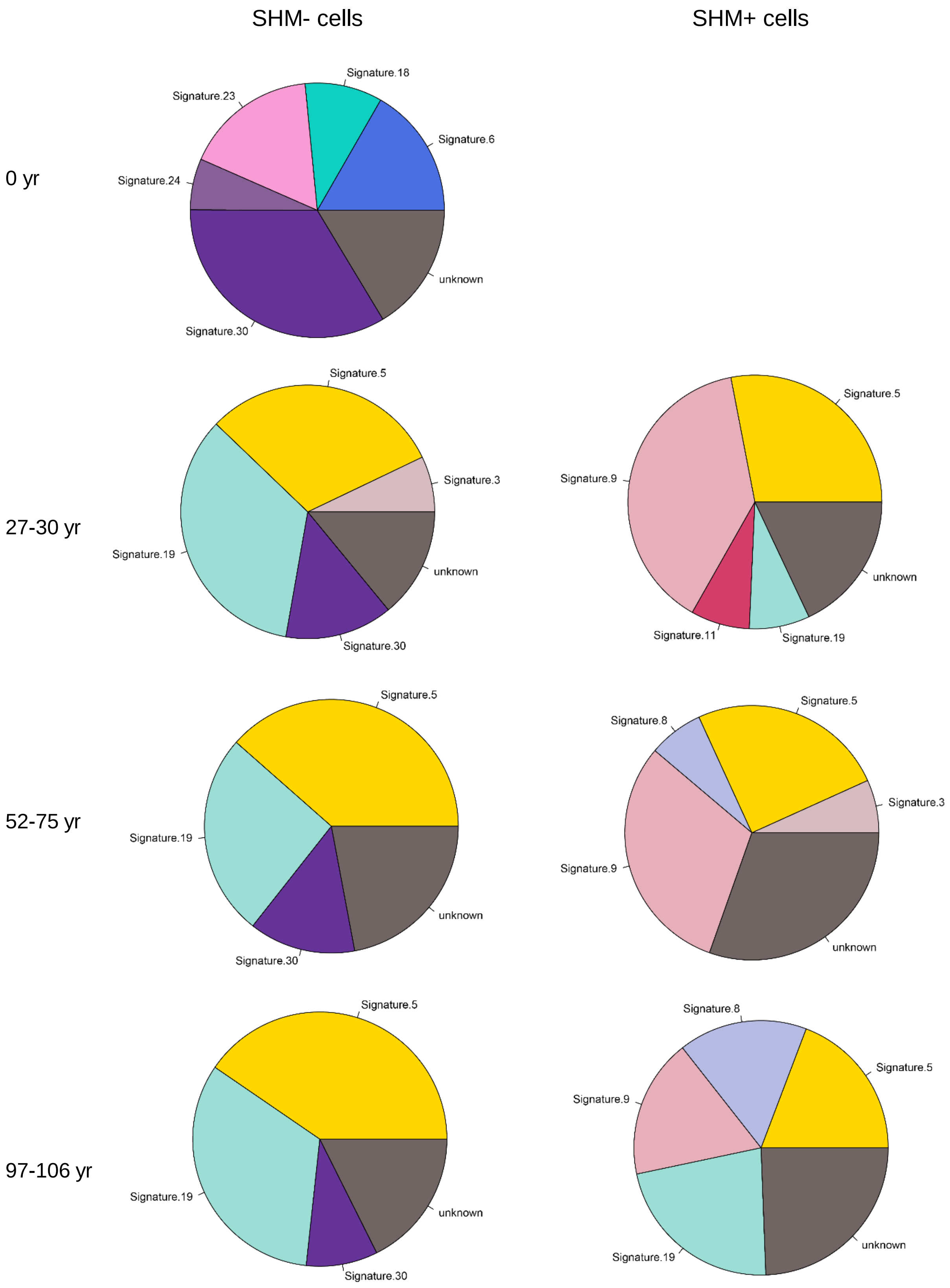


**Supplementary Fig. 13 | Mutation signatures by refitting to the 30 COSMIC signatures.**

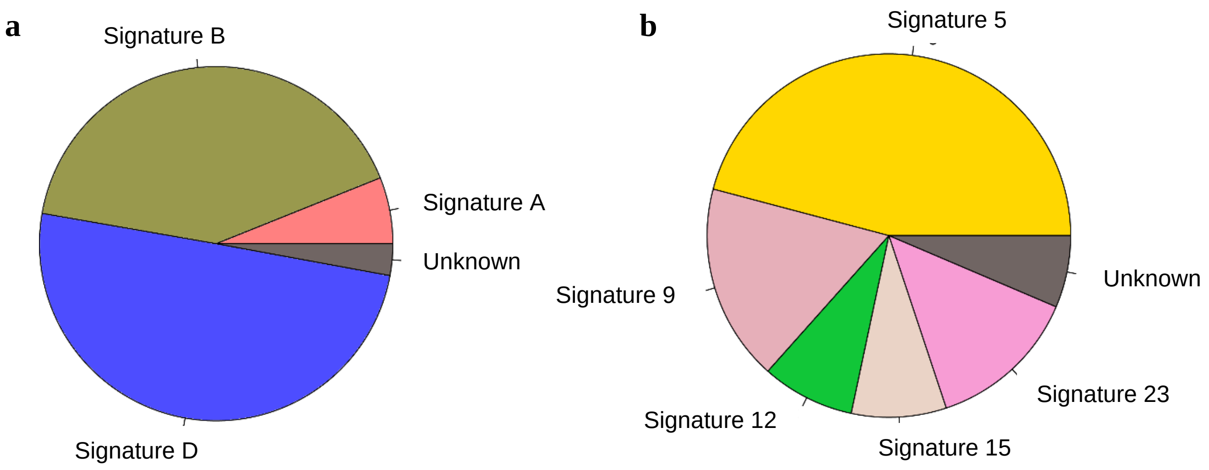


**Supplementary Fig. 14 | Mutation signature at the 24 hotspot regions.**
**a.** The analysis by refitting to the four signatures identified in this study. **b.** The analysis by refitting to the 30 COSMIC signatures.

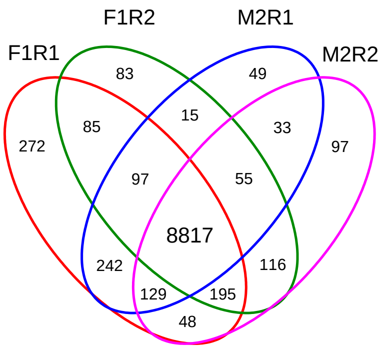


**Supplementary Fig. 15 | Replications of RNA sequencing.**
A Venn diagram shows the overlap of transcribed genes between four RNA sequencing replicates.

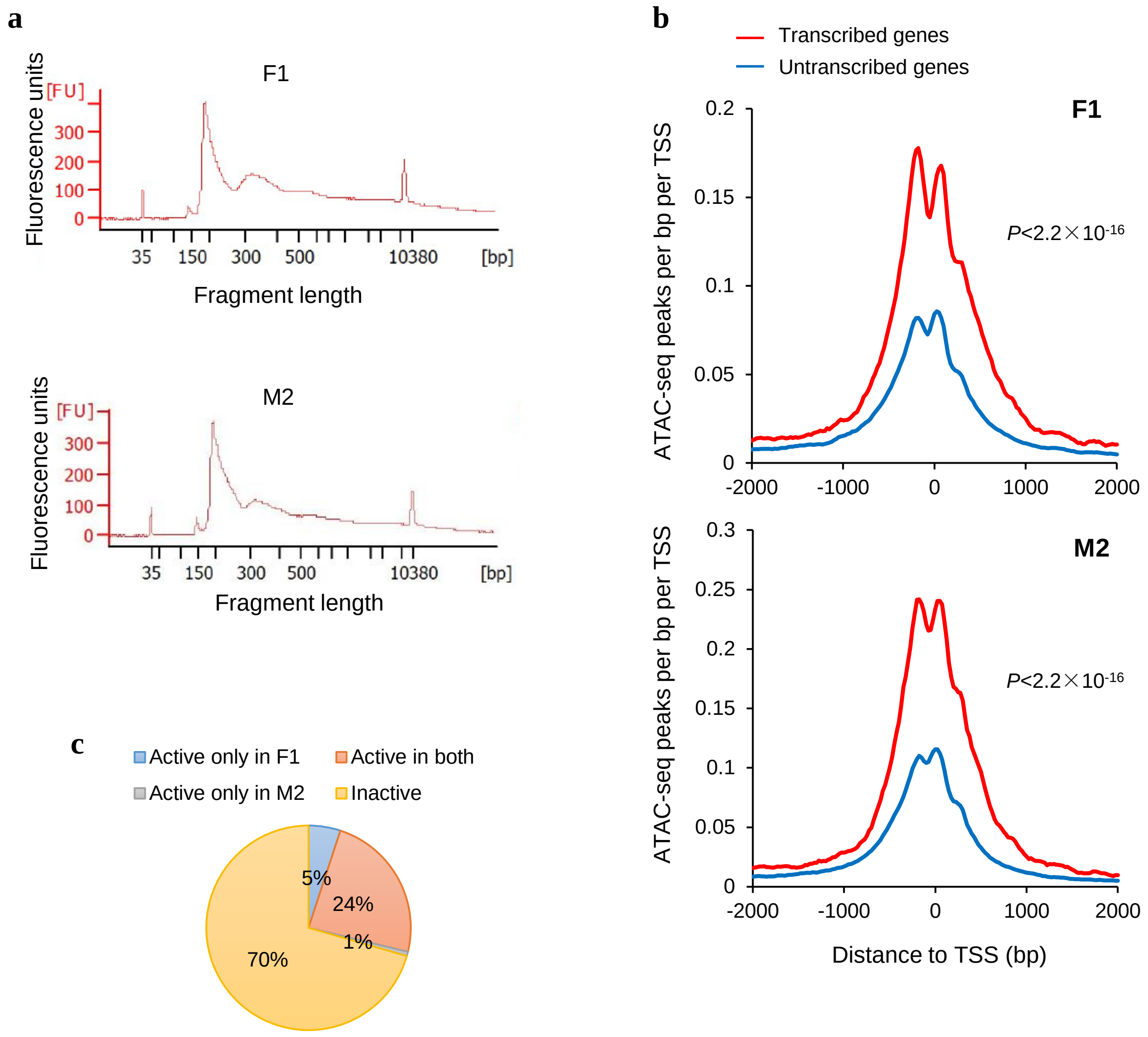


**Supplementary Fig. 16 | ATAC sequencing.**
**a.** Bioanalyzer of ATAC sequencing library. **b.** ATAC sequencing peaks are significantly enriched in active genes than inactive genes in both of our ATAC sequencing samples (both P<2.2×10-16, Wilcoxon signed-rank test, one tailed). **c.** Fraction of active TF binding regions identified using ATAC sequencing from two replicates, F1 and M2. The total TF binding regions were identified by the ENCODE from multiple cell types.
